# Erythrocyte signalling is critical for *Plasmodium falciparum* invasion

**DOI:** 10.1101/2022.07.18.500419

**Authors:** James Jia Ming Yong, Xiaohong Gao, Prem Prakash, Soak Kuan Lai, Ming Wei Chen, Jason Jun Long Neo, Julien Lescar, Hoi Yeung Li, Peter R. Preiser

## Abstract

Successful *Plasmodium falciparum* merozoite invasion requires the activation of red blood cell (RBC) signalling pathways. The binding of parasite ligand reticulocyte binding protein homologue 5 (RH5) to its host receptor Basigin is essential for merozoite invasion and triggers a Ca^2+^ influx in RBCs. Here we observed that RH5-bound RBCs form a multimeric protein complex containing Basigin, CD44 and β2-adrenergic receptor (β2AR), suggesting that RH5-Basigin interaction is functionally associated with the host cAMP signalling pathway. Interestingly, we detected a characteristic rise in cAMP levels in the RBC upon RH5-Basigin interaction, which can be blocked by G protein and cAMP-synthesising adenylyl cyclase (AC) inhibitors. Furthermore, we demonstrated that RBC L-type Ca^2+^ channel inhibitor and cAMP signalling inhibitors are able to block merozoite invasion. Checkerboard invasion inhibition assay containing different combinations of signalling inhibitors also exhibited a drastic amplification of inhibition levels, indicating that these signalling proteins are functioning in a common signalling cascade to activate the L-type Ca^2+^ channels. Taken together, this study provides new insights into the role of a host cAMP-Ca^2+^ signalling pathway during merozoite invasion and sheds new light on antimalarial therapeutic strategies to tackle the high infection rate and growing threat of drug resistant parasites.

**Key Points:**

1. A pre-existing Basigin-associated membrane protein complex undergoes increased protein assembly upon RH5 binding on the RBC surface.
2. *Plasmodium falciparum* merozoite exploits host cAMP signalling to initiate Ca^2+^ influx in the RBC.

## Introduction

Malaria is a major burden in developing countries today, resulting in more than 600,000 deaths in 2020. *Plasmodium falciparum* has been responsible for most severe malaria cases and malaria-related mortalities ^1^. Successful *P. falciparum* invasion requires a network of protein-protein interactions between the host and parasite ^2,3^. The interaction between parasite ligand reticulocyte-binding protein homolog 5 (RH5) and host receptor Basigin is essential for merozoite invasion and triggers RBC Ca^2+^ influx ^4–7^.

While this host-parasite relationship has garnered significant attention, our current knowledge of RBC signalling events during merozoite invasion is still limited. It is though clear that the parasites have manipulated specific host cell signalling mechanisms via RH5-Basigin interaction to trigger RBC Ca^2+^ signalling and propagate downstream phosphorylation on cytoskeletal proteins to promote the rearrangement of the RBC cytoskeleton, which is critical for merozoite insertion and tight junction formation ^4,7^. Recent evidence has indicated that the cytoplasmic domain of Basigin is non-functional and does not directly activate downstream effectors upon RH5-Basigin binding ^8^, suggesting that RH5-bound Basigin relays signals from an extracellular to intracellular environment via a host cell signalling cascade.

Interestingly, G protein-coupled receptor (GPCR) activation of β-adrenergic receptor (βAR) by stimulatory G protein α (Gαs) modulates RBC membrane rigidity ^9,10^. Gαs-stimulated GPCR subsequently activates adenylyl cyclase (AC) to catalyse the cyclization of ATP into cAMP ^11,12^. Furthermore, β2AR agonists reportedly stimulate AC-mediated cAMP production in RBCs and lead to increased *P. falciparum* infection that can be perturbed by β2AR antagonists and Gα inhibitory peptides ^13,14^, indicating that GPCR-mediated activation of RBC cAMP signalling is critical for invasion. Elevated levels of intracellular cAMP will then interact with cAMP-dependant protein kinase A (PKA) regulatory subunits (PKAr), releasing PKA catalytic subunits (PKAc) that phosphorylate the serine and threonine residues on its effector proteins ^15–17^. These PKA-sensitive substrates include RBC cytoskeleton proteins, such as adducin and protein 4.1R ^18,19^, and potentially trigger the destabilisation of the cytoskeletal network in line with our previous findings on the significant loss of ankyrin and adducin from the RBC cytoskeleton and increase in the RBC spectrin mesh size upon RH5 binding ^4^. It is also notable that while PKA reportedly activates voltage-gated Ca^2+^ channel in nucleated cells ^20,21^, it is still unclear whether it is responsible for the activation of host Ca^2+^ voltage-gated channels and Ca^2+^-sensitive kinases to trigger the phosphorylation of RBC cytoskeleton proteins during invasion.

Here we provided evidence of a multimeric membrane protein complex on RH5-bound RBCs that undergoes increased assembly of RBC membrane proteins including CD44 and β2AR, and detected a rise in RBC cAMP level that drives RBC Ca^2+^ influx upon RH5-Basigin interaction. We demonstrated that cAMP signalling proteins and L-type Ca^2+^ channels are involved in RBC Ca^2+^ influx, suggesting an interplay between host cAMP and Ca^2+^ signalling that is required for invasion. Taken together, RH5-Basigin interaction potentially triggers an activation cascade of RBC membrane proteins in a multimeric complex to activate host cAMP-Ca^2+^ signalling that directs the invasion process.

## Methods

### Ethics statement

The whole blood was donated by healthy non-malarial immune adult volunteers either at the National University Hospital, Singapore or Fullerton Healthcare at Nanyang Technological University. Informed consent given was written. Informed consents were obtained from all donors in accordance with protocols approved by Institutional Review Board of Nanyang Technological University, Singapore (IRB-2018-02-031, IRB-2019--09-047, IRB-2019--09-042 and IRB-2020--11-047).

### *Plasmodium falciparum* parasite culture

Parasite strain 3D7 (MR4) were cultured in fresh RBCs as described ^22^.

### Isolation of *Plasmodium falciparum* merozoites

Mature schizonts were harvested as described ^23^ and treated with 10 μM E64 (Sigma). Pellet containing E64-treated schizonts were resuspended in cRPMI medium and filtered through a 2 µm Isopore polycarbonate membrane (Sigma) to purify free merozoites.

### Generation of RBC ghosts

Fresh RBC pellets were split into two sets. One set is kept on ice. The other set is incubated with 0.2 mg/ml rRH5 in RPMI medium. Two sets of pellets were lysed in 1x hypotonic lysis buffer comprising of 1.43 mM Na_2_PO4 and 5.7 mM Na_2_HPO_4_ to form RBC ghost, followed by solubilised in IP lysis buffer (Thermo Scientific). Solubilised RH5-bound RBC ghosts were incubated with Ni-NTA beads and further resuspended in 500 mM imidazole to elute the rRH5-bound proteins.

### Western blot analysis of RBC ghosts

Unbound RBC ghosts were separated on 10% SDS-PAGE. RH5-bound and unbound RBC ghosts were separated on 5% Native-PAGE. The gels were transferred onto nitrocellulose membranes (0.2 μm) (Bio-Rad) and probed with Basigin, CD44 and β2AR antibodies (Abcam) at 1:2,000 dilution, respectively, followed by HRP-linked secondary antibodies (1:5,000) and enhanced chemiluminescence (GE healthcare). α-His antibody (Qiagen) was used at 1:5,000 dilution to detect the presence of rRH5 in Native-PAGE.

### Generation of resealed RBC containing membrane-impermeable signalling inhibitors

Fresh RBCs were lysed and resealed as described previously ^14^. During lysis, membrane-impermeable inhibitors (Table 1) were added, respectively. Influenza Hemagglutinin (HA) peptide (Sigma) was added into a separate set as a control.

**Table 1.** Properties of signalling inhibitors used in the *P. falciparum* merozoite invasion inhibition assays and RBC cAMP-Ca^2+^ signalling assays.

| Permeability | Compounds | Abbreviation | Company | Target | References |
| --- | --- | --- | --- | --- | --- |
| Membrane-permeable | BIM46187 | - | MedChem Express | G protein | 95 |
|  | 2',5'-Dideoxy-adenosine | 2',5'-ddA | Sigma | AC | 96 |
|  | PKA (14-22) | - | Sigma | PKAc | 97 |
|  | Verapamil | - | Sigma | L-type Ca<sup>2+</sup> channels | 98 |
| Membrane-impermeable | G Protein antagonist peptide | GPA | Sigma | βAR | 33,99 |
|  | NF449 | - | Santa Cruz Biotechnology | Gαs | 100 |
|  | 2'-Deoxy-adenosine 3'-monophosphate | 2'-d-3'-AMP | Santa Cruz Biotechnology | AC | 101 |
|  | Rp-8-OH-cAMPS | - | Santa Cruz Biotechnology | PKAr | 102 |
|  | PKA (6-22) | - | Sigma | PKAc | 103,104 |
|  | N-methyl verapamil | N-methyl Vera | Abcam | L-type Ca<sup>2+</sup> channels | 105 |

### cAMP immunoassay

RBC cAMP dynamics were measured using the cAMP Direct Immunoassay Kit (Abcam) following Abcam’s protocol. 10 µL RBCs were incubated in the absence and presence of rRH5 protein (0.2 mg/ml) in a total volume of 100 µL in 1x PBS and measured at 0, 2, 4, 6, 8, and 10 min post-incubation. 50 μM Forskolin (Abcam) was used as a positive control. 80 μM BIM46187 and 80 μM 2’,5’-ddA (Table 1) were applied to RBCs in the presence of rRH5, respectively, and measured after 2-min incubation. rRBCs were prepared in the absence and presence of GPA, NF449 and 2’-d-3’-AMP, respectively. rRBCs loaded with HA peptide were prepared as an internal control. cAMP level was measured as described in rRBCs in the presence of rRH5 after 2-min incubation. The fold change in cAMP level in the RBC or rRBC was compared to that in RBC only or rRBC only.

### Merozoite invasion inhibition assay with membrane-impermeable inhibitors

Membrane-impermeable inhibitors (Table 1) was loaded in rRBCs at 2-fold increasing concentrations from 0.1953125 μM to 100 μM. Purified merozoites were added at 2% hematocrit. rRBCs pre-loaded in the absence and presence of 100 μM HA peptide were used as a positive control and an internal positive control, respectively. Invasion control experiment was performed in normal RBCs in the absence and presence of membrane-impermeable inhibitors (Table 1). HA peptide was added to normal RBCs as an internal control. Invasion inhibition assays were performed and (%) invasion inhibition was calculated as described ^24^.

### Visualisation and measurement of RBC Ca^2+^ by fluorescent microscopy and fluorescence plate reader

Fluo-4 AM-labelled (Invitrogen) RBCs were incubated with rRH5 protein at 0.2 mg/ml in the absence and presence of membrane-permeable inhibitors including 80 μM BIM46187, 80 μM 2’,5’-ddA, 40 μM PKA (14-22) and 40 μM verapamil, respectively. The culture was applied under Nikon Eclipse Ti Inverted Microscope. 20× objective lens was used to observe and capture a wider field of RBC. The above-mentioned samples were prepared for a real-time RBC Ca^2+^ measurement as described ^24^. Preloaded rRBCs were incubated with either freshly purified merozoites or rRH5 at 0.2 mg/ml in the absence and presence of membrane-impermeable inhibitors (Table 1) at 100 μM, respectively. rRBCs loaded with HA peptide (100 μM) was used as an internal positive control. Time-resolved changes of cytosolic Ca^2+^ levels inside RBCs and rRBCs were assessed using a fluorescence plate reader (Infinite M200; Tecan) and analysed as described ^24^.

### Live video microscopy

Live imaging of merozoite invasion into Fluo-4AM labelled rRBCs was performed as described ^4^ to analyse the effect of membrane-impermeable inhibitors (Table 1) at 100 μM on Ca^2+^ signal. Freshly prepared labelled rRBCs were used as a positive control. Labelled rRBCs preloaded with 100 μM HA peptide were prepared as an internal positive control. Mature schizonts were incubated with compound-loaded rRBCs and observed under Zeiss LSM710 confocal microscope. Images were analysed by using Zen black software (Carl Zeiss).

### Checkerboard invasion inhibition assay

Purified merozoites were incubated with rRBCs pre-loaded with the combinations of impermeable inhibitors (Table 1) with N-methyl Vera at 2% hematocrit. Purified merozoites incubated with rRBCs were used as positive control. Invasion inhibition was measured as described ^24^. Invasion in the presence of inhibitors was compared with positive controls of invasion as described above. The Minimum Inhibitory Concentration (MIC) and the Fractional Inhibitory Concentration (FIC) index were used to calculate the isobologram in Figure 6 ^25,26^.

### Statistical analysis

Statistical comparison was done using one-way ANOVA as appropriate. A *p-*value of <0.05 was considered statistically significant. The *p*-value is provided in individual experiment if required.

## Results

### RH5-bound Basigin forms a multimeric protein complex containing CD44 and β2-adrenergic receptor on the RBC membrane

RH5-Basigin interaction triggers RBC Ca^2+^ influx that is critical for *P. falciparum* merozoite invasion ^4,5,27^. Interestingly, Basigin associates with β2AR via a mediator protein in a pre-existing membrane complex, which triggers specific signalling events to facilitate meningococcal invasion ^28^. Functional interaction between Basigin and CD44 on the membrane during invasion further suggests that CD44 functions as a binding partner of RH5-bound Basigin ^29^. To confirm the detection of Basigin, β2AR and CD44 on RBC ghosts, we extracted RBC ghosts and performed western blot analysis. All three antibodies (Abs), α-Basigin, α-β2AR and α-CD44, recognised their expected proteins in size respectively (Figure 1A), confirming the presence of Basigin, β2AR and CD44 on the RBC surface. To address whether RH5-Basigin binding forms a complex with CD44 and β2AR, we have successfully purified full-length recombinant RH5 protein with His-tag (rRH5) in an expected size of 63 kDa (Supplemental Figure 1A) as described ^4^. Properly folded rRH5 directly binds to RBC in the erythrocyte binding assay, indicating that rRH5 acts as native RH5 (Supplemental Figure 1B). We further extracted RBC ghosts in the absence and presence of functional rRH5 and carried out Native-PAGE and western blot. A band at approximately 300 to 400 kDa in size was recognised by α-Basigin, α-CD44 and α-β2AR Abs in the RBC ghost extraction, respectively (Figure 1B), indicating a pre-existing Basigin-associated complex is formed on the RBC membrane. Strikingly, all three antibodies detected a large increase in protein complex size to approximately 800 kDa upon rRH5 binding (Figure 1B), demonstrating that RH5-Basigin interaction leads to an increased assembly of diverse RBC membrane proteins such as CD44 and β2AR. α-His antibody was used as a loading control of rRH5 and confirmed the presence of rRH5 in the extraction. The presence of β2AR in a multimeric RH5-bound RBC complex therefore highlights that RH5-bound Basigin may kickstart host GPCR-mediated signalling pathways to facilitate RBC Ca^2+^ influx.

**Figure 1:**
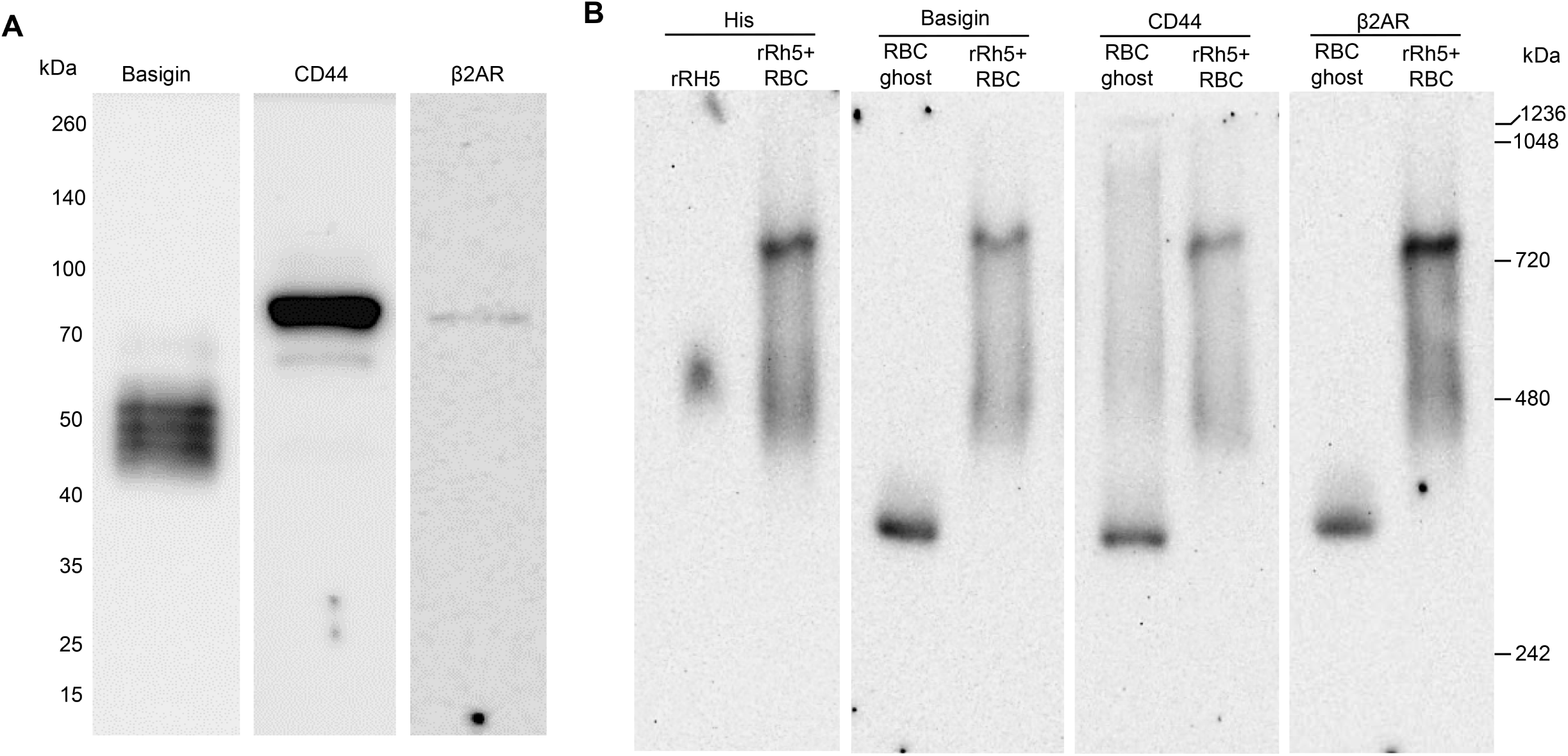
Basigin associates with membrane proteins to form a multimeric complex on the RBC membrane. (A) Unbound RBC ghosts were separated on a single 10% SDS-PAGE and probed with Basigin, CD44 and β2AR antibodies. (B) Basigin is part of RBC membrane complex containing CD44 and β2AR. Unbound RBC ghosts (RBC ghost) and eluted samples of RH5-bound RBC ghosts (rRH5+RBC) were separated on a single 5% Native-PAGE and probed with Basigin, CD44 and β2AR antibodies. Anti-His antibody (His) was also used to detect rRH5 itself (rRH5) and RH5-bound RBC ghosts (rRH5+RBC). The protein molecular weight (kDa) is indicated on either the left or right side.

### RH5-Basigin interaction triggers a rise in host cAMP level to facilitate RBC Ca^2+^ influx

Interactions between the cAMP and Ca^2+^ signalling pathways occur at multiple stages in mammalian cells to regulate signalling activities, and in turn modulate the degree of cellular response ^30,31^. To address whether RH5-Basigin binding triggers host cAMP signalling, we measured the dynamic changes of RBC cAMP levels for 10 min with 2-min intervals upon rRH5 binding using a cAMP competitive ELISA. The RBC cAMP level in the absence of rRH5 was used as a negative control. AC activator Forskolin ^32,33^ was used as a positive control. Incubation of RBC with Forskolin elevated intracellular cAMP levels in a time-independent manner (Figure 2A). A stronger rise in cAMP levels to a peak of 10.63 μM at about 2-min time point, was observed upon the addition of recombinant RH5 showing that RH5 triggers the increase of intracellular cAMP levels in RBCs. The cAMP level decreased to 5.27 μM at 10 min (Figure 2A). The observed increase in cAMP upon rRH5 binding can be blocked by G protein inhibitor BIM46187 and AC inhibitor 2’,5’-ddA (Table 1) (Figure 2B), confirming that RH5-Basigin interaction triggers RBC cAMP generation. To examine whether an elevated cAMP level regulates the Ca^2+^ uptake into the RBC, direct visualisation of induced RBC Ca^2+^ increase was performed using fluorescent microscopy. Meanwhile, accumulative Ca^2+^ signals were obtained at 600 s by TECAN fluorescent plate reader. Fluo-4 AM labelled RBC alone was used as a negative control with only few cells fluorescing (Figure 2C-i). Consistent with our previous findings, rRH5 triggered an increase of RBC cytosolic Ca^2+^ that causes the whole RBC to fluoresce (Fig Figure 2C-ii) and this is inhibited by BIM46187 and 2’,5’-ddA (Figure 2C-iii and iv; Figure 2D, bar chart in green). Moreover, PKA inhibitor PKA (14-22) and L-type Ca^2+^ channel inhibitor verapamil (Table 1) dramatically reduced the RBC Ca^2+^ signal in the presence of rRH5 (Figure 2C-v and vi; Figure 2D, bar chart in blue). Our findings clearly indicate that elevated cAMP levels induce an increase of RBC cytosolic Ca^2+^ level that enable Ca^2+^ uptake through the host L-type Ca^2+^ channels. Overall, these data are consistent with RH5 binding to Basigin alone being sufficient to promote Ca^2+^ uptake through Ca^2+^ transporters into the RBC via changes in RBC cAMP level during invasion.

**Figure 2:**
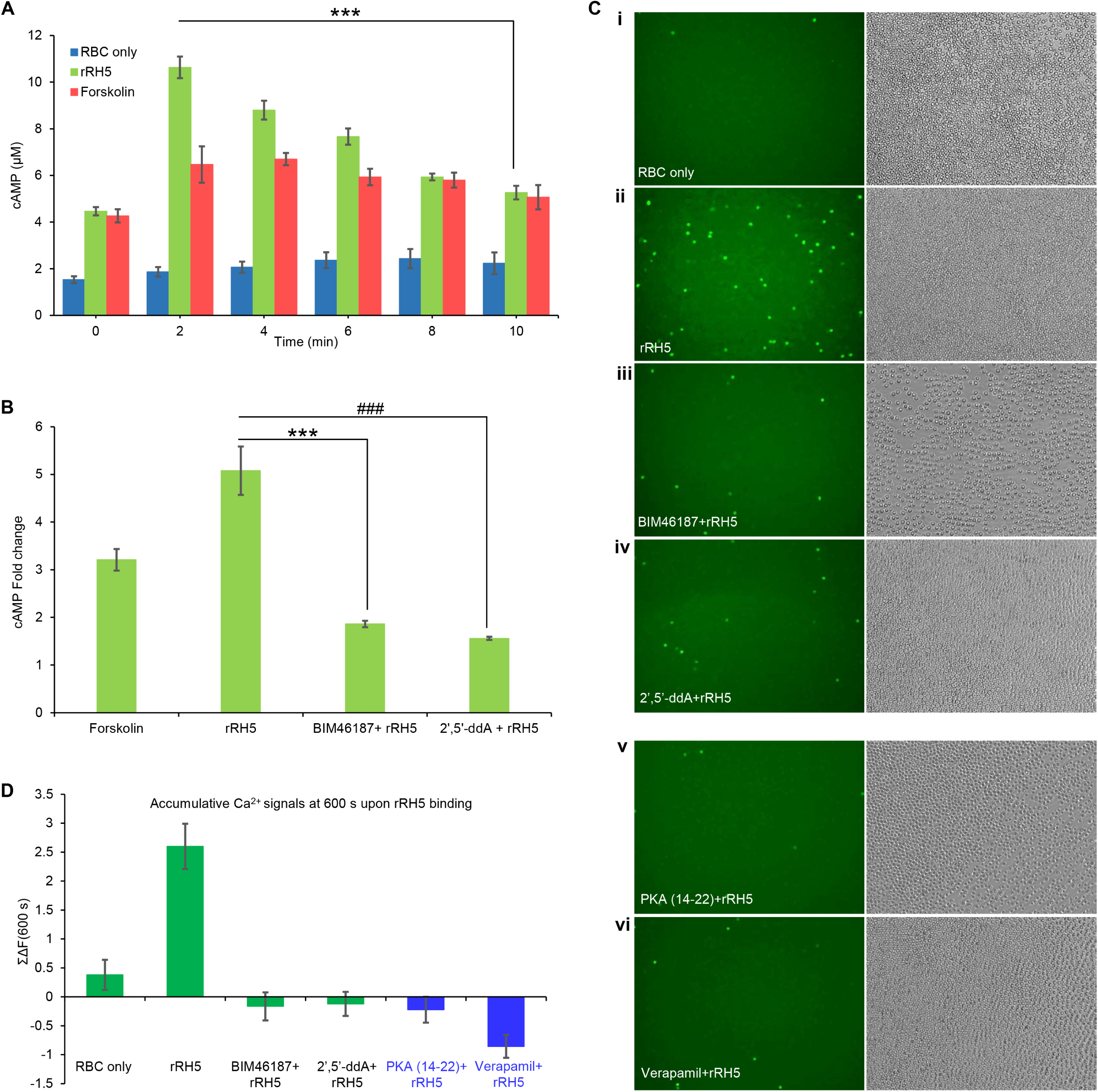
Recombinant RH5 (rRH5) triggers a rise in cytosolic cAMP level that promotes RBC Ca^2+^ influx. (A) cAMP immunoassay of RBCs. The dynamic changes of RBC cAMP level were measured in the absence (RBC only) and presence of 0.2 mg/mL recombinant RH5 protein (RH5) within 10 min in 2-min intervals. Forskolin was used as a positive control (Forskolin). Experimental data presented as mean ± s.e.m, n=3. \*\*\**p* = 0.00060379 indicates the significant difference on the cAMP level at 2 min compared to that at 10 min. (B) cAMP inhibition immunoassay of RBCs. The dynamics of RBC cAMP level was measured upon rRH5 binding for 2 min in the absence (RH5) and presence of G protein inhibitor BIM46187 (BIM46187+RH5) or adenylyl cyclase (AC) inhibitor 2′,5′-ddA (2′,5′-ddA+RH5). Forskolin was used as a positive control (Forskolin). The cAMP reading in RBC alone was used as a negative control. The fold change in cAMP level in the samples was compared to that in the negative control and plotted as a bar chart. \*\*\**p* = 0.003262353 and ###*p* = 0.002275767 indicate the significant difference on fold change in RBC cAMP level upon rHR5 binding between in the absence (rRH5) and presence of BIM46187 (BIM46187+RH5) or 2′,5′-ddA (2′,5′-ddA+RH5), respectively. Experimental data presented as mean ± s.e.m, n=3. (C) Fluorescence images of RBC Ca^2+^. Fresh RBCs labelled with Fluo-4AM were incubated in the absence (RBC only) and presence of rRH5 (rRH5) and presence of BIM46187 (BIM46187+RH5), 2′,5′-ddA (2′,5′-ddA+RH5) or PKA (14-22) (PKA (14-22)+rRH5) prior to rRH5 binding for 10 min. Fluorescence (left) and bright field (right) images were taken under 20× objective lens. (D) RBC Ca^2+^ measurement by using fluorescence plate reader. The Ca^2+^ signal intensity of the samples obtained from (**c**) were also measured. Dynamics of RBC cytosolic Ca^2+^ level was measured over 600 s by fluorescence plate reader. ΣΔF(t) which reflects the cumulative change in RBC cytosolic Ca^2+^ was plotted as a bar chart. Experimental data presented as mean ± s.e.m, n=3.

### Host cAMP signalling proteins and L-type Ca^2+^ channels are essential for merozoite invasion

RBCs possess cAMP-dependent signalling pathways that regulate biomechanical properties of the RBC membrane to adapt to extracellular signals in the blood microenvironment ^34–36^. Since RH5 binding to Basigin is a prerequisite for cAMP-mediated Ca^2+^ signalling pathway, cAMP signalling proteins are expected to be involved in RBC signalling during invasion. To prevent off-target inhibition of parasite-encoded cAMP signalling proteins ^37^, merozoite-based experiments were performing by preloading RBCs with each membrane-impermeable inhibitor of interest (Table 1), using a modified protocol from Murphy et al ^14^ to form “Resealed RBC” (rRBC). Influenza Hemagglutinin (HA) peptide was loaded into rRBCs as an internal control and shown by dot blot assay using anti-HA antibody to be specifically captured within these rRBCs (Supplemental Figure 2A), while no signal was detected in empty rRBCs. Importantly, rRBCs are invaded with equal efficiency as normal RBCs (Supplemental Figure 2B, C), highlighting that rRBCs are suitable for merozoite invasion inhibition studies. To assess whether the presence of these membrane-impermeable inhibitors in the culture media had any effect, merozoite invasion assays were carried out using normal RBCs in the presence of impermeable cAMP signalling inhibitors and L-type Ca^2+^ channel inhibitor (Table 1). None of the inhibitors elicited a significant inhibitory effect on invasion as compared to the control without inhibitors (Supplemental Figure 2D), confirming the suitability of these inhibitors to investigate the role of RBC signalling molecules during invasion.

We evaluated the impact of preloading the different inhibitors into rRBC on either cAMP levels or Ca^2+^ signalling using rRH5. While both Empty rRBCs and HA-loaded rRBCs showed a significant increase in cAMP levels upon incubation with rRH5, this was drastically blocked by βAR inhibitor GPA, G protein inhibitor NF449 and AC inhibitor 2’-d-3’-AMP (Figure 3A), respectively, providing further evidence that the binding of RH5 to Basigin promotes host cAMP signalling. Similarly, rRBCs pre-loaded with both Fluo-4AM and different inhibitors showed a significant reduction of Ca^2+^ signalling in the presence of GPA, NF449 and 2’-d-3’-AMP as compared to the Empty rRBCs and HA-loaded rRBCs controls (Figure 3B), confirming an RH5-induced cAMP regulated signalling event. The inclusion of PKAr inhibitor Rp-8-OH-cAMPS, PKAc inhibitor PKA (6-22) and L-type Ca^2+^ channel inhibitor N-methyl verapamil showed a significant reduction in Ca^2+^ signalling consistent with our previous data using normal RBCs, and further confirming that RH5-Basigin interaction manipulate host cAMP signalling to promote RBC Ca^2+^ influx through the host L-type Ca^2+^ channels.

**Figure 3:**
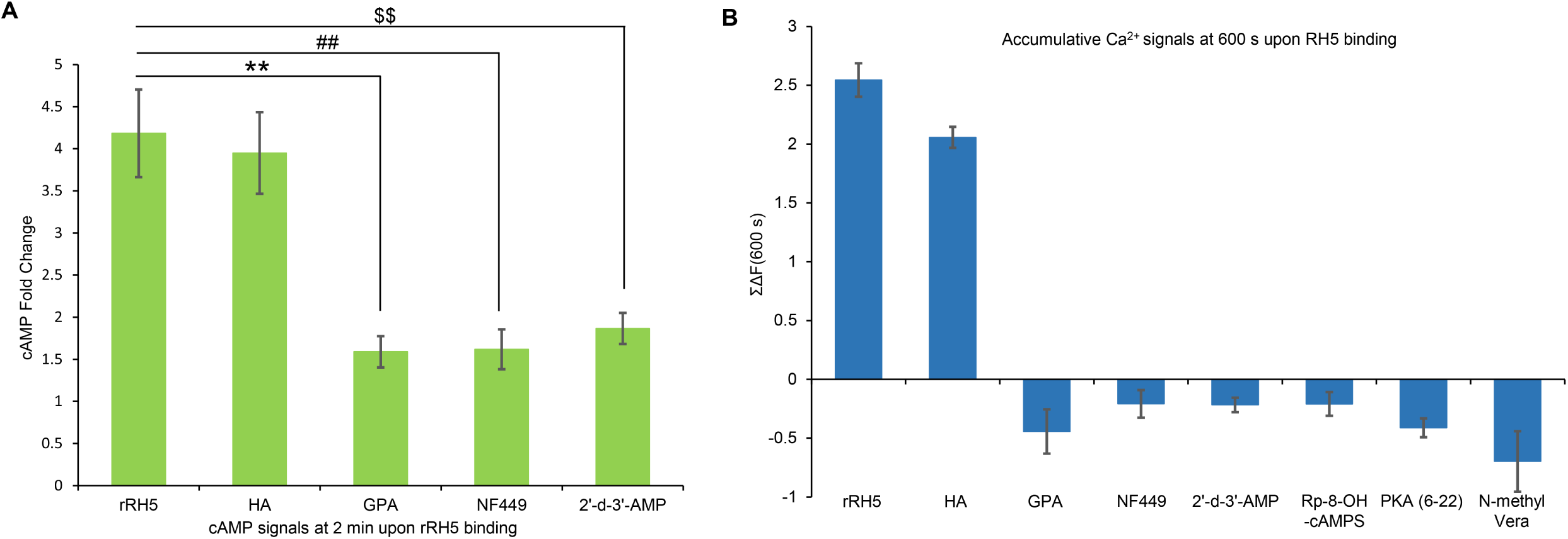
RH5-Basigin interaction drives RBC Ca^2+^ influx via cAMP signalling in resealed RBCs. (A) cAMP inhibition immunoassay of rRBCs. The dynamics of RBC cAMP level was measured upon rRH5 binding for 2 min in the absence (rRBC) and presence of βAR inhibitor GPA, G protein inhibitor NF449 or adenylyl cyclase inhibitor 2’-d-3’-AMP. The cAMP reading in HA-loaded rRBCs was used as an internal positive control (HA). \*\**p* = 0.010384776, ##*p* =0.012368378 and $$*p* =0.015778993 indicate the significant difference on fold change in rRBC cAMP level upon rHR5 binding between in the absence (rRH5) and presence of GPA (GPA+rRH5), NF449 (NF449+rRH5) or 2’-d-3’-AMP (2’-d-3’-AMP+rRH5), respectively. Experimental data presented as mean ± s.e.m, n=3. (B) rRBC Ca^2+^ measurement by using fluorescence plate reader. Fluo-4AM labelled resealed RBCs were incubated with rRH5 protein in the absence (rRH5) and presence of inhibitors GPA, NF449, 2’-d-3’-AMP, Rp-8-OH-cAMPS, PKA (6-22) or N-methyl verapamil (N-methyl Vera), respectively. HA-loaded resealed RBC (HA) was used as an internal positive control. Dynamics of RBC cytosolic Ca^2+^ level was measured over 600 s by fluorescence plate reader. ΣΔF(t) which reflects the cumulative change in RBC cytosolic Ca^2+^ was plotted as a bar chart. Experimental data presented as mean ± s.e.m, n=3.

Having shown that the membrane-impermeable inhibitors significantly impact the cAMP induced Ca^2+^ signalling cascade, we evaluated whether they are essential for merozoite invasion. Invasion assays using rRBCs preloaded with impermeable inhibitors (Table 1) showed dose-dependent invasion inhibition for each inhibitor tested (Figure 4) as compared to HA-loaded rRBC control. The data confirms that the host cAMP signalling proteins and L-type Ca^2+^ channels are essential for invasion and take part in cAMP-Ca^2+^ signalling transduction during invasion. Interestingly, cAMP-sensitive PKA is reportedly involved in the activation of L-type Ca^2+^ voltage-gated channels ^20^, and these Ca^2+^ channels can regulate intracellular Ca^2+^ levels in the RBC by enabling Ca^2+^ uptake from the extracellular space into the cytosol ^38,39^. These data strongly imply the involvement of a cAMP-dependent signalling pathway induced by RH5-Basigin interaction that triggers the increase of RBC cytosolic Ca^2+^ level during invasion.

**Figure 4:**
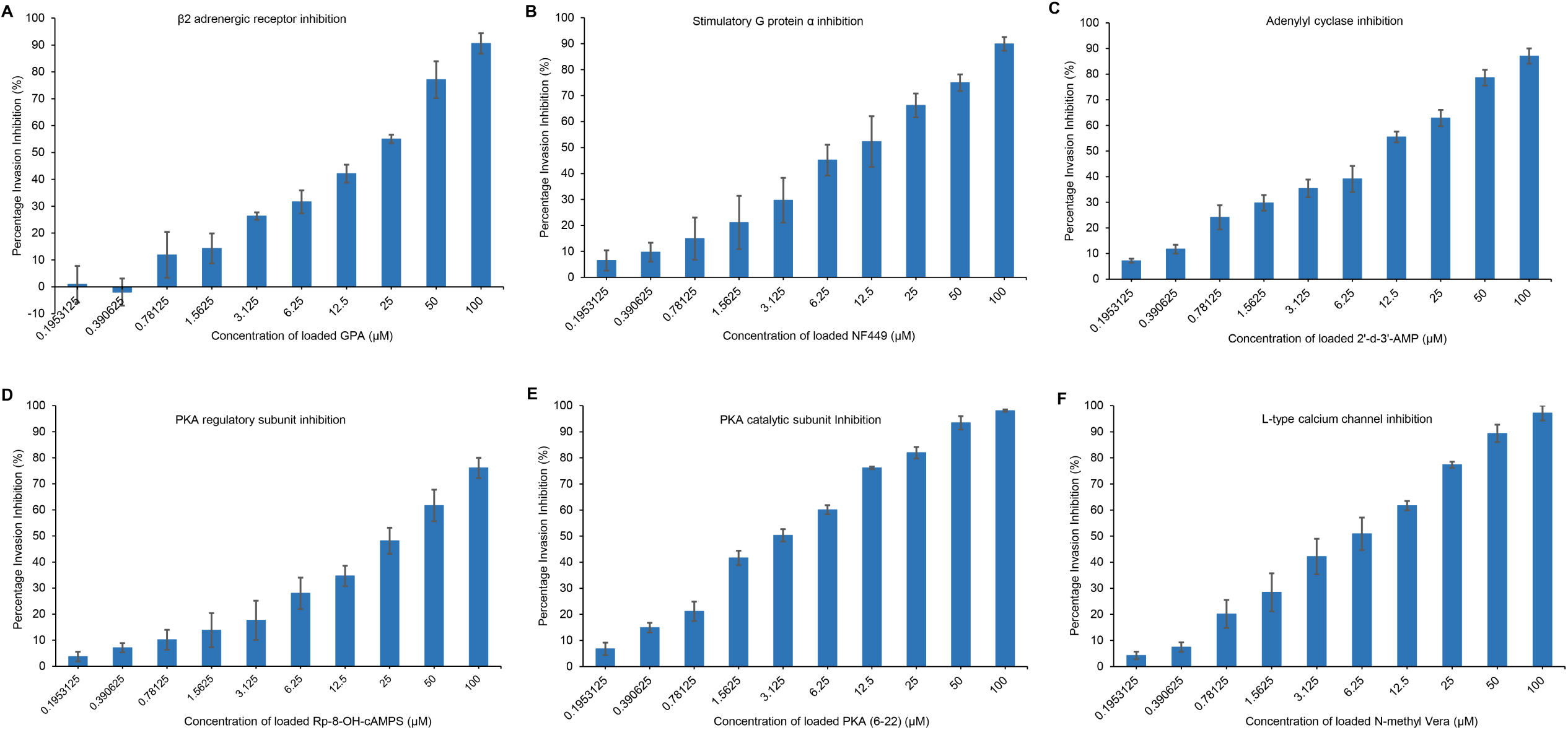
cAMP-Ca^2+,^ signalling inhibitors block merozoite invasion. Merozoite invasion inhibition assay was performed using purified merozoites with resealed RBCs that were preloaded with impermeable inhibitors such as (A) GPA, (B) NF449, (C) 2’-d-3’-AMP, (D) Rp-8-OH-cAMPS, (E) PKA (6-22) and (F) N-methyl verapamil (N-methyl Vera), respectively, at 2-fold increasing concentrations from 0.1953125 μM to 100 μM. Resealed RBCs preloaded with HA peptide were used as a positive control. Bar chart represents the percentage invasion inhibition of 3D7 merozoite that was compared with the invasion into resealed RBCs preloaded with HA peptide in the absence of inhibitors. Experimental data were reported as mean ± s.e.m; n=3.

### Host cAMP signalling proteins and L-type Ca^2+^ channels induce RBC Ca^2+,^ influx during merozoite invasion

The tightly regulated Ca^2+^ influx from the extracellular space into the RBC cytosol triggered by RH5-Basigin interaction is required for invasion ^4^. Since the cAMP signalling inhibitors and L-type Ca^2+^ channel inhibitor block invasion, we investigated whether there is a clear relationship between RBC Ca^2+^ signalling and invasion inhibition. In line with previous observations, live video microscopy of merozoite invasion using rRBCs preloaded with Fluo-4AM, in the absence and presence of HA peptide (Figure. 5A-B left panels, Supplemental Video 1, 2), exhibited successful invasions that begin with a Ca^2+^ influx inside the rRBC after merozoite initial attachment and reorientation, and the signal rapidly spread throughout the entire rRBC. These invaded rRBCs subsequently undergo extensive deformation and Ca^2+^ decrease until the signal disappeared after 90 s. Moreover, rRBC Ca^2+^ influx during invasion showed a significant increase in fluorescence intensity when compared to uninfected rRBCs in the vicinity, as shown in Figure 5A-B right panels. To assess the impact of cAMP signalling inhibitors and L-type Ca^2+^ channel inhibitor on Ca^2+^ signalling in rRBCs during invasion, we loaded each inhibitor into rRBCs before staining with Fluo-4AM for detection using live video microscopy. Interestingly, in all inhibitor-loaded rRBCs, Ca^2+^ flux was blocked after the initial merozoite attachment (Figure 5C-5H and Supplemental Video 3-8), indicating that the cAMP signalling proteins and L-type Ca^2+^ channels are directly responsible for RBC Ca^2+^ uptake, which is required for merozoite invasion. Our data provide further evidence that *P. falciparum* parasites can exploit the host cAMP signalling pathway downstream of RH5-Basigin interaction to promote the activation of Ca^2+^ voltage-gated channels, which in turn enable RBC Ca^2+^ influx to facilitate invasion. The inhibitory effects of cAMP signalling inhibitors and L-type Ca^2+^ channel inhibitor on rRBC Ca^2+^ were further quantified as described ^4,24^. Merozoite invasion leads to a significant increase in detectable Ca^2+^ signals in positive controls, but the signals were significantly reduced in inhibitor-loaded rRBCs (Figure 5I), confirming that the cAMP signalling proteins and L-type Ca^2+^ channels play key roles in regulating host Ca^2+^ signalling during invasion. In line with previous observations, the results signify that RH5-Basigin interaction triggers cAMP and Ca^2+^ signalling in the RBC to facilitate successful invasion.

**Figure 5:**
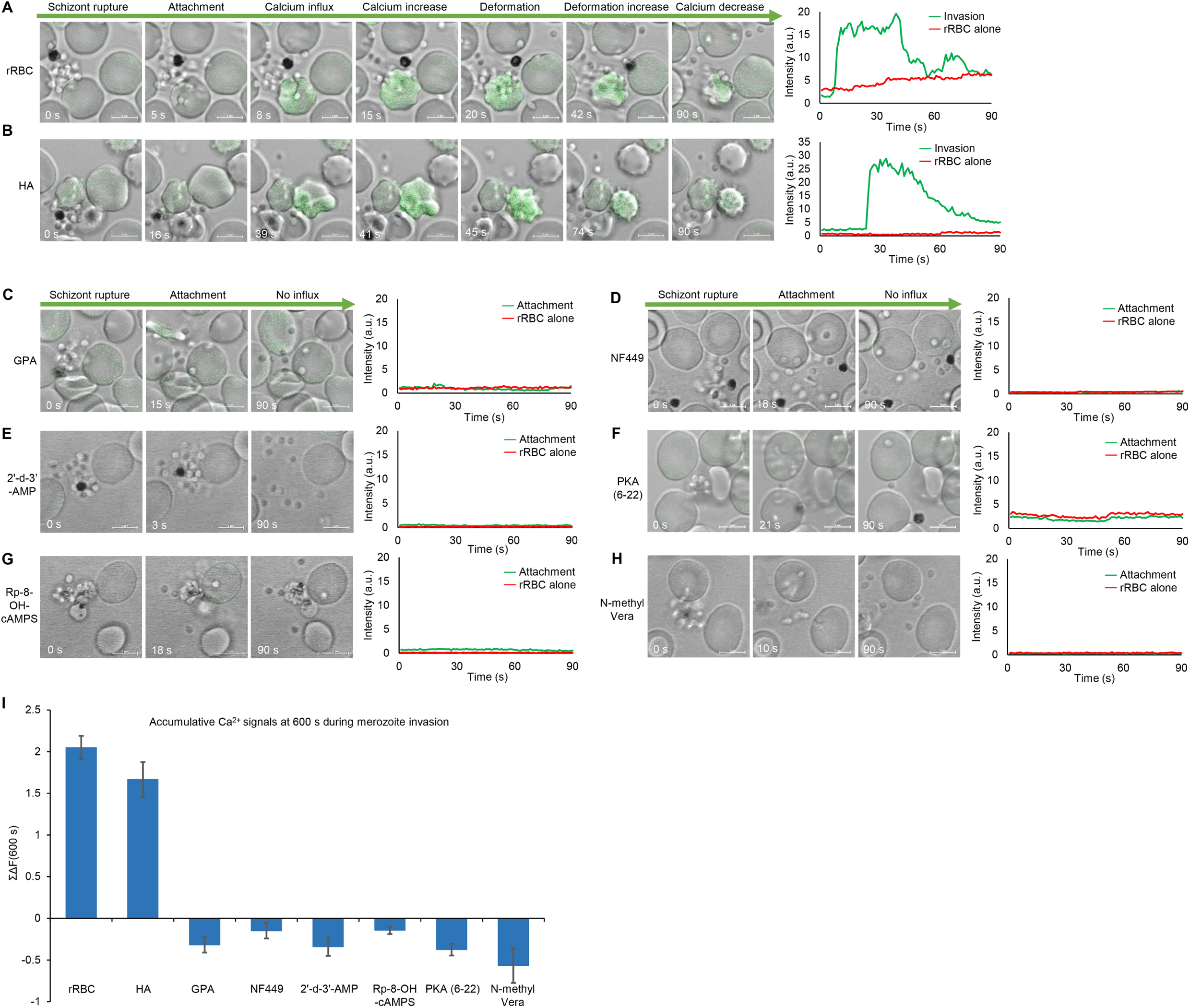
Live video microscopy of merozoite invasion into resealed RBCs. Representative snapshots taken from time-lapse live video microscopy of 3D7 merozoites invasion in the Fluo-4AM labelled resealed RBCs in the absence (rRBC) (A) and presence of HA peptide (HA) (B) Merozoite attaches to the RBC after the schizont rupture and reorients followed by Ca^2+^ influx that spreads throughout the entire resealed RBC. The Ca^2+^ influx causes a massive deformation, and the signal starts to decrease when echinocytosis happens (Supplementary Video 1-2). Representative video snapshots were also taken in the Fluo-4AM labelled resealed RBCs in the presence of (C) GPA, (D) NF449, (E) 2’-d-3’-AMP, (F) Rp-8-OH-cAMPS, (G) PKA (6-22) and (H) N-methyl verapamil (N-methyl Vera), respectively. In the presence of (C-H), the merozoite released from a mature schizont attaches and apically reorients, but it is unable to penetrate the RBC with no Ca^2+^ influx after 90 s (Supplementary video 3-8). Dynamics of rRBC Ca^2+^ signals was measured. Ca^2+^ fluorescence intensity was indicated in green line ((Invasion) in (A) and (B)) and ((Attachment) in (C)-(H)) compared with uninfected rRBCs (rRBC alone in red line) within 90 s. Time was shown in seconds. Scale bar = 5 μm. (I) rRBC Ca^2+^ measurement by using fluorescence plate reader during merozoite invasion. Fluo-4AM labelled rRBCs were incubated with 3D7 merozoites in the absence (rRBC) and presence of inhibitors GPA, NF449, 2’-d-3’-AMP, Rp-8-OH-cAMPS, PKA (6-22) or N-methyl verapamil (N-methyl Vera), respectively. HA-loaded resealed RBC (HA) was used as an internal positive control. Dynamics of RBC cytosolic Ca^2+^ level was measured over 600 s by fluorescence plate reader. ΣΔF(t) which reflects the cumulative change in RBC cytosolic Ca^2+^ was plotted as a bar chart. Experimental data presented as mean ± s.e.m, n=3.

### Host cAMP signalling proteins and L-type Ca^2+^, channels function synergistically during merozoite invasion

The synergistic effect of cAMP and Ca^2+^ signalling is known to regulate key physiological functions in mammalian cells ^30,31,35^. As RBC Ca^2+^ uptake is potentially driven by the activation of Ca^2+^ voltage-gated channels, we aim to determine whether a crosstalk between the cAMP and Ca^2+^ signalling pathways influences the entry of extracellular Ca^2+^ signals via the L-type Ca^2+^ channels during invasion. We performed a checkerboard invasion inhibition assay to measure the combined inhibition level of rRBCs loaded with dual combinations of impermeable signalling inhibitors. The impermeable L-type Ca^2+^ channel inhibitor N-methyl verapamil was selected as the reference inhibitor to be paired with GPA, NF449, 2’-d-3’-AMP, Rp-8-OH-cAMP, or PKA (6-22), respectively. Interestingly, all dual inhibitor combinations exhibited drastically higher invasion inhibition levels when compared to that of individual inhibitor (Figure 6A-i-v, Supplemental Figure 3). This is evident when the inhibitor concentrations were increased, the invasion inhibition level of paired checkerboard squares peaked higher than the expected sum of individual inhibition levels by both inhibitors (Supplemental Figure 3), suggesting a synergistic relationship between host cAMP and Ca^2+^ signalling that is functionally active in merozoite invasion. Furthermore, an isobologram of all five checkerboard invasion inhibition assays was applied between N-methyl verapamil and its paired cAMP signalling inhibitors. Since the Fractional Inhibitory Concentration (FIC) index <0.5 indicates a synergistic inhibitory interaction ^25,26^, our isobolographic analysis revealed potent synergism in invasion inhibition by N-methyl verapamil and its paired cAMP signalling inhibitor in all five dual inhibitor combinations (Figure 6B), which is consistent with our previous findings. Our data further highlight that a host-specific crosstalk exists between the cAMP and Ca^2+^ signalling pathways during invasion. Taken together, we provide a body of evidence that the host cAMP signalling proteins and L-type Ca^2+^ channels are functioning synergistically in a GPCR-mediated activation of RBC cAMP-Ca^2+^ signalling cascade that regulates RBC Ca^2+^ influx during merozoite invasion.

**Figure 6:**
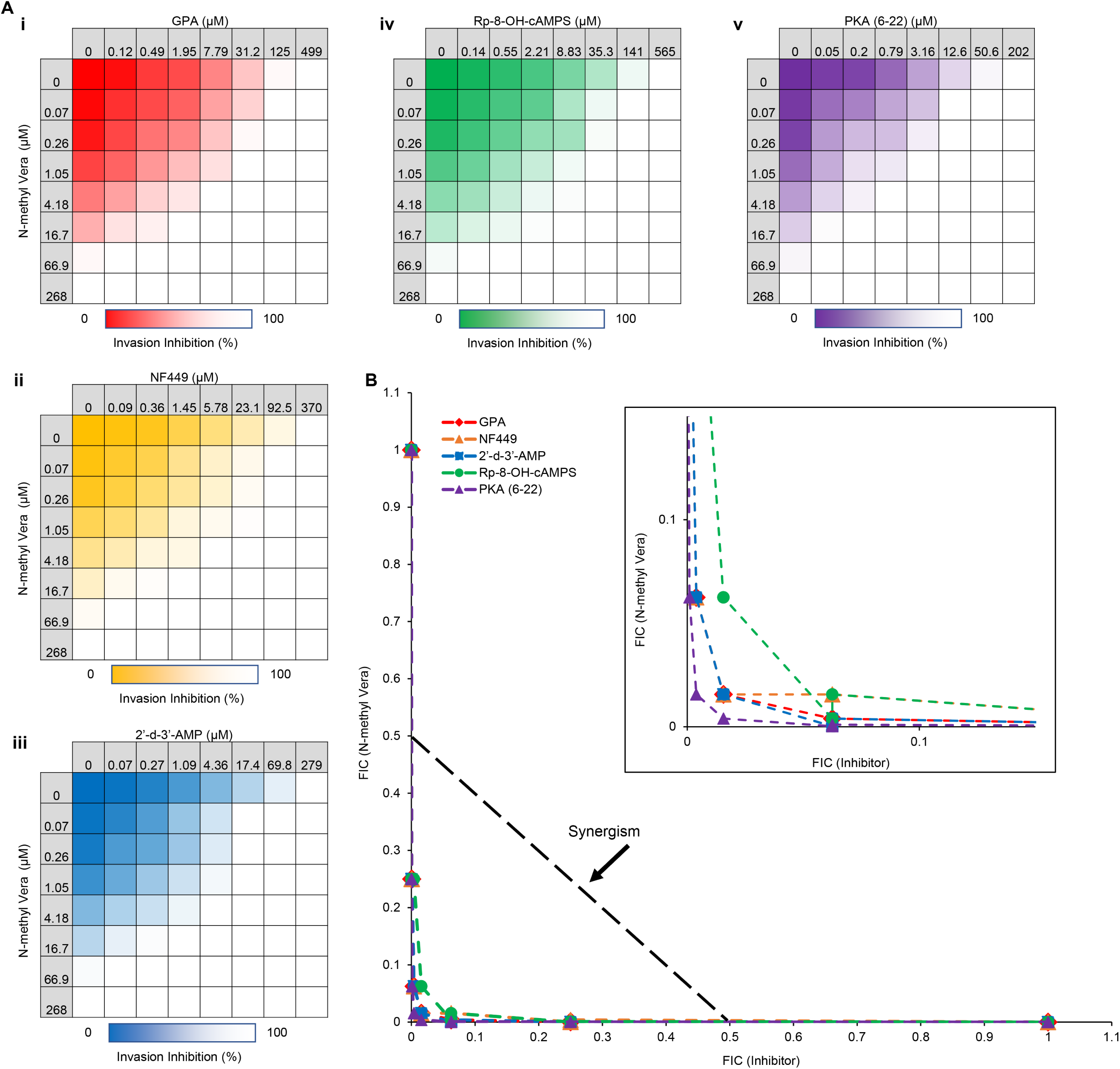
Synergistic effect of cAMP-Ca^2+^ signalling inhibitors during merozoite invasion. (A) Inhibition heat plots of 3D7 merozoite invasion into rRBCs. Resealed RBCs were pre-loaded with dual combinations of Ca^2+^ channel blocker (N-methyl Vera) and other cAMP signalling inhibitors including (i) GPA (red), (ii) NF449 (yellow), (iii) 2’-d-3’-AMP (blue), (iv) Rp-8-OH-cAMPS (green), and (v) PKA (6-22) (purple). All inhibitors were loaded in rRBCs at about 4-fold increasing concentrations including N-methyl Vera shown in each table vertically, while the other inhibitors shown horizontally in colour matrices. Colours in the dose-response matrices from the darkest to white indicate different levels of invasion inhibitions from 0% to 100% obtained from Supplemental Figure 3. (B) Isobologram plot of five different checkerboard invasion inhibition data from Supplemental Figure 3. The Minimum Inhibitory Concentration (MIC) of each inhibitor in the checkerboard invasion inhibition assay was calculated by using the equation MIC = MIC [(A in combination) / MIC (A alone)], in which [MIC (A alone)] is the Minimum Inhibitory Concentration to achieve complete invasion inhibition by inhibitor A alone, while [MIC (A in combination)] is the concentration of inhibitor A that exhibit the Minimum Inhibitory Concentration to achieve complete invasion inhibition when paired with another inhibitor. To illustrate the type of inhibitory interaction in each dual inhibitor combination, all five checkerboard data were graphically represented with an isobologram. The isobologram curve was formed by plotting the MIC values of varying inhibitor concentrations tested in the checkerboard assay. The Fractional Inhibitory Concentration (FIC) index <0.5 indicates a synergistic interaction, while the FIC index in the range of 0.5 to 0.9 indicates a partially synergistic interaction. Additionally, the FIC index in the range of 1.0 to 2.0 indicates an additive effect, whereas the FIC index >4.0 indicates an antagonistic effect. The range of Fractional Inhibitory Concentration (FIC) values surrounding the black dotted line with arrow in black from 0 to 0.5 represents a synergistic isobole. Data points below this line indicate potent synergism, while the data points above this line indicate partial synergism. Inset: zoomed plot at 0.15 of both x-and y-axis for better visualisation.

## Discussion

RH5-Basigin interaction can trigger RBC Ca^2+^ influx that potentially leads to host Ca^2+^-dependent phosphorylation events to regulate the RBC cytoskeletal remodelling ^4,24^, suggesting that the RBC, which was once thought to be passive, plays a key role in facilitating merozoite invasion. This implies that the parasite has manipulated specific host signalling pathways downstream of RH5-Basigin interaction to activate voltage-gated Ca^2+^ channels during invasion. Additionally, Basigin can associate with β2AR, a member of the GPCR protein family that mediates the activation of signalling pathways including cAMP, in a pre-existing membrane complex to initiate host cell signalling events ^10,13,14,28^. Furthermore, Basigin has a propensity to form complexes with multiple membrane-associated proteins including cyclophilins and matrix metalloproteinases (MMP), which are key players in signalling transduction and tumour invasion ^40,41^.

Basigin knockdown in differentiating erythrocytes from hematopoietic stem cells blocks merozoite invasion, indicating that Basigin is required for *P. falciparum* invasion ^5^. Furthermore, a CD44 knockout JK-1 cell line enhances anti-Basigin–dependent inhibition of invasion, signifying that Basigin and CD44 form a functional association during merozoite invasion ^29^. Moreover, reticulocytes expressing a truncated Basigin protein lacking the C-terminal cytoplasmic domain exhibited similar invasive susceptibility to that of unmodified reticulocytes, indicating that C-terminal of Basigin is not essential for invasion ^8^ and signalling. Consistent with this our data demonstrates that Basigin forms a pre-existing membrane complex with CD44 and β2AR prior to invasion and potentially recruits additional RBC membrane proteins upon RH5 binding, suggesting that the increased protein assembly on the RBC membrane is critical in driving downstream invasion events.

Our study reveals new insights into a potential crosstalk between RBC cAMP and Ca^2+^ signalling during invasion. RH5-Basigin interaction induces a rise in cytosolic cAMP levels and this in turn triggers RBC Ca^2+^ influx. Our findings indicate that the convergence between the cAMP and Ca^2+^ signalling pathways results in the release of PKA catalytic subunits that may directly or indirectly activate voltage-gated L-type Ca^2+^ channels ^20,21,42^ to promote Ca^2+^ uptake from the extracellular space into the RBC cytosol. While there has been mounting evidence of PKA phosphorylation sites on the α and β subunits of L-type Ca^2+^ channels that increase channel activities ^20,21,42^, it remains unclear whether RH5-mediated cAMP signalling triggers PKA-mediated phosphorylation of L-type Ca^2+^ channels thereby upregulating channel function. It is though clear that cAMP signalling plays an upstream regulatory role in triggering RBC Ca^2+^ uptake, and the perturbation of host cAMP signalling protein activities effectively blocks this Ca^2+^ influx, and ultimately, merozoite invasion.

Entry of external Ca^2+^ into RBCs is required for Ca^2+^-dependent activities because the RBC lacks organelles, such as mitochondria that commonly serve as intracellular Ca^2+^ source in mammalian cells ^38^. The RBC tightly controls the strong Ca^2+^ concentration gradient between the cytosol and extracellular environment by maintaining low basal Ca^2+^ concentrations in the cytosol with its low Ca^2+^ membrane permeability and membranous Ca^2+^ transporters ^43^. The presence of L-type Ca^2+^ channels on the RBC membrane therefore enables a regulated route of Ca^2+^ transport from the extracellular space into the RBC cytosol. This is in line with previous evidence that the removal of Ca^2+^ from the culture media using EGTA blocked RH5-mediated Ca^2+^ influx ^4^. Interestingly, we observed that after initial merozoite attachment, Ca^2+^ influx in the RBC spreads throughout the cell, indicative of the entry of extracellular Ca^2+^ through voltage-gated Ca^2+^ channels that are distributed along the RBC membrane.

Taken together, we have identified a critical RBC signalling cascade that is activated by the RH5-Basigin interaction and shown that these host signalling proteins can be targeted to block invasion. We propose that upon RH5 binding on the RBC surface, a pre-existing Basigin-associated membrane protein complex containing CD44 and other unknown host proteins undergoes increased protein assembly and recruitment that subsequently activates the GPCRs such as β-adrenergic receptor (Figure 7-I). The activation of β-adrenergic receptor in turn triggers the host cAMP-PKA signalling pathway that either directly or indirectly activates Ca^2+^ channel to enable RBC Ca^2+^ influx during invasion (Figure 7-II-V). The activation of PKA also potentially results in the phosphorylation of RBC cytoskeletal proteins, thereby destabilising the cytoskeletal network and loosening the RBC membrane to promote parasite entry (Figure 7-V). Our current findings provide new evidence on the ability of the malaria parasite to manipulate host cell derived cAMP and Ca^2+^ signalling pathways in invasion and highlights the importance of focusing on host derived signalling molecules as new intervention targets.

**Figure 7:**
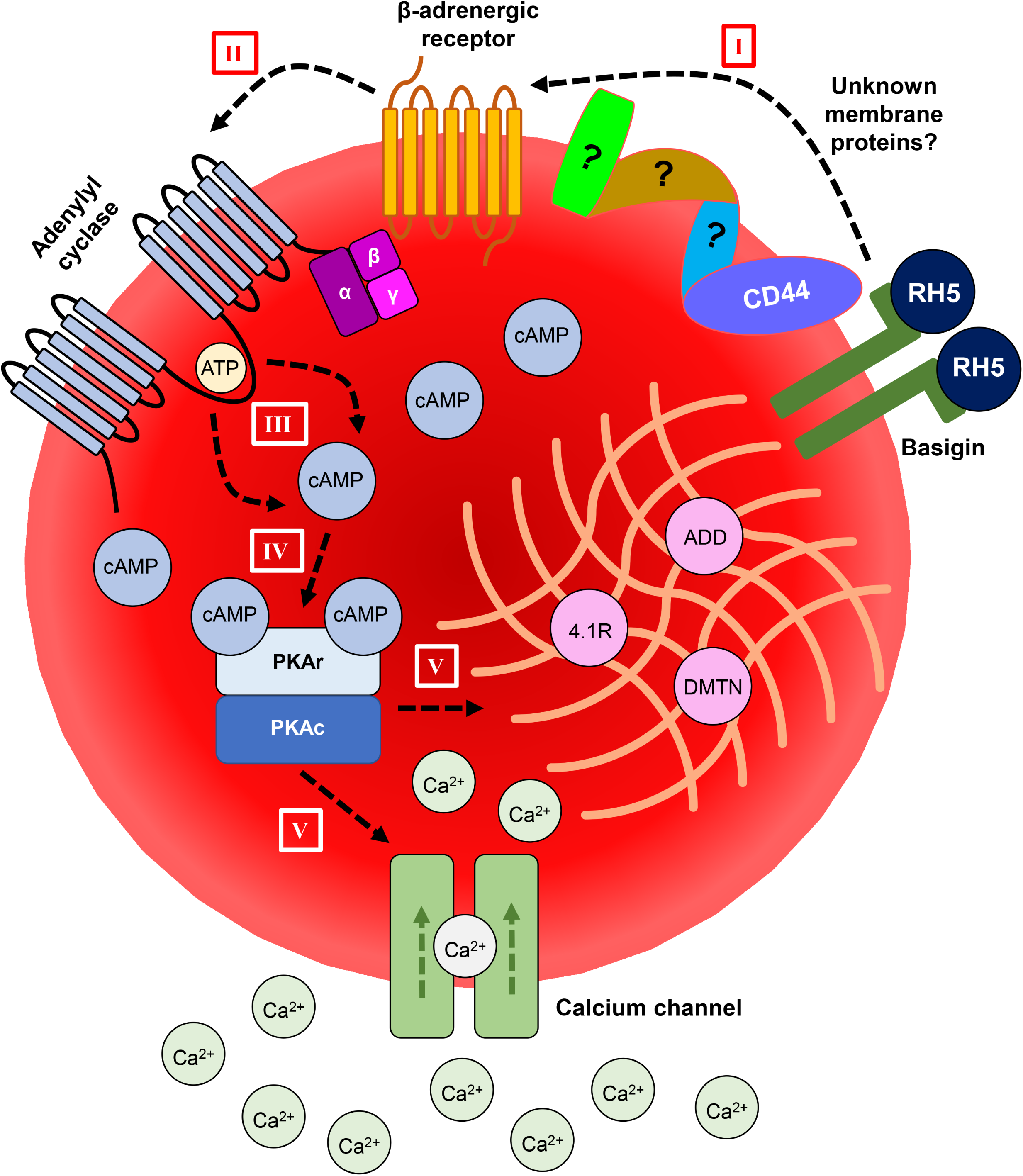
Schematic model of RBC cAMP-Ca^2+^, signalling during merozoite invasion. RBC cAMP-Ca^2+^ signalling is manipulated by the parasite during merozoite invasion. RH5-Basigin interaction triggers an activation cascade of neighbouring molecules around Basigin such as CD44 and other unknown RBC proteins, which activates the β-adrenergic receptors (I). Activation of βAR triggers the dissociation of stimulatory G protein alpha (Gαs) that binds to the transmembrane adenylyl cyclase (II). Activation of adenylyl cyclase (AC) generates cAMP from ATP via 3’-5’-ATP-cyclization reaction (III). cAMP molecules bind to the PKA (IV) and subsequently activates the kinase. PKA may directly or indirectly be involved in the activation of L-type Ca^2^, channels. Alternatively, PKA may also phosphorylate the cytoskeleton proteins such as adducin and erythrocyte membrane protein band 4.1R, to promote overall changes of RBC cytoskeleton structure (V).

## Acknowledgements

We are grateful to all the blood donors. We also thank the NTU Protein Production Platform for expression and purification of recombinant RH5. This research is supported by Singapore Ministry of Education Academic Research Fund Tier 2 (MOE2017-T2-1-034 and MOE-T2EP30121-0013). The funders had no role in study design, data collection and analysis, decision to publish, or preparation of the manuscript.

## Author contributions

J.J.M.Y and X.G contributed extensively and equally to the experiments, analysed the data and wrote the manuscript. J.J.M.Y, X.G, P.P and P.R.P conceived the experiments. M.W.C, J.J.L.N and L.J performed the recombinant RH5 protein expression and purification. J.J.M.Y, X.G, S.K.L and H.Y.L performed the live video microscopy and analysed the data. P.R.P supervised the project, analysed the data, and wrote the manuscript.

